# Mendelian randomization identifies the potential causal impact of dietary patterns on circulating blood metabolites

**DOI:** 10.1101/2020.10.09.332924

**Authors:** Nele Taba, Hanna-Kristel Valge, Andres Metspalu, Tõnu Esko, James F. Wilson, Krista Fischer, Nicola Pirastu

## Abstract

Nutrition plays an important role in the development and progress of several health conditions, but the exact mechanism is often still unclear. Blood metabolites are likely candidates to be mediating these relationships, as their levels are strongly dependent on the frequency of consumption of several foods/drinks. Understanding the causal effect of food on metabolites is thus of extreme importance. To establish these effects we utilized Two-sample Mendelian randomization using the genetic variants associated with dietary traits as instrumental variables. The estimates of single-nucleotide polymorphisms’ effects on exposures were obtained from a recent genome-wide association study (GWAS) of 25 individual and 15 principal-component dietary traits, whereas the ones for outcomes were obtained from a GWAS of 123 blood metabolites measured by nuclear magnetic resonance spectroscopy. We identified 417 potentially causal links between food and metabolites, replicating previous findings, such as the association between increased oily fish consumption and higher DHA, and highlighting several novel associations. Most of the associations were related to very-low-density, intermediate-density (IDL) and low-density lipoproteins (LDL). For example, we found that constituents of IDL particles and large LDL particles were raised by coffee and alcohol while lowered by an overall healthier diet and fruit consumption. Our results represent one of the first examples of the estimates of long-term causal effects of diet on metabolites and start bridging the gap in the mechanistic understanding linking food consumption to its health consequences.

## Introduction

Nutrition plays an important role in the development and progress of several diseases, such as obesity [1], type II diabetes (T2D) [2], cardiovascular diseases (CVD) [3–5] and cancer [6,7]. These in turn create a high burden for individuals, society, the economy and health-care and thus prevention is of great importance. In many cases the mechanism by which food consumption acts on health is still unclear. Blood metabolites are promising candidates for filling this gap. Metabolites have been shown to be important in the onset of a wide range of diseases such as type II diabetes [8,9], incident cardiovascular events [10,11], dementia [12,13] and colorectal cancer [14,15], and indicative of mortality [16], and are thus likely mediators for at least some of the food-health relationships.

In recent years, the progress in quantifying metabolites has allowed investigation of the relationship between food and blood or urine metabolite levels. Studies have shown associations between metabolomic profile and intake of fruit and vegetables [17], coffee [14], alcohol [18] and a wide range of dietary patterns. A more detailed overview of the current state of the field is summarized by Guasch-Ferré, Bhupathiraju and Hu [19] and Brennan and Hu [20]. Most studies in this field are observational and are thus limited by the typical biases which affect nutritional epidemiology (i.e. reporting bias, strong correlation between the studied variables etc.) and are therefore unfit to detect causal relationships. Moreover, even in feeding studies conducted under very strict and controlled conditions, the effects could be measured only on a short-term basis and on limited sample size, which in turn limits statistical power.

Nevertheless, the feeding studies provide convincing evidence that dietary intake has causal effects on the metabolic profile, highlighting the potential to assess dietary intake via investigation of metabolomic profiles. Such studies have been applied to: the percentage of dietary intake coming from carbohydrates/fat/protein [21], the effects of low-glycaemic index diet [22] and orange juice on proline betaine [23]. Change in biomarkers can be caused by consumption or non-consumption of various dietary items and therefore variations in blood metabolites are likely candidates to be mediating the effects of food on health. Thus, detecting causal relationships between dietary choices and biomarkers might reveal more insight into the mechanism by which food affects health.

A possible alternative to approaches of observational- and feeding studies is the method of Mendelian Randomization (MR) [24]. MR exploits the natural randomization of the alleles associated with exposure in the population to measure the long term effects of the trait on the outcome of interest. MR assumes that people with the allele increasing the exposure will be exposed to this difference for their whole life and thus by comparing the effect of the allele on the exposure and the outcome it is possible to derive the effect of the exposure on the outcome. MR relies on the results coming from genome-wide association studies (GWAS) which are generally publicly available and does not require the direct involvement of the participants. It is thus extremely cost-effective and it is possible to apply it to contexts where randomized controlled trials would be unethical (for example alcohol consumption). If carefully conducted, MR is exempt from the biases that are typical of observational studies. MR has been successfully used in many different contexts including nutritional epidemiology in the cases, for example, of milk [25,26], alcohol [27,28], and coffee consumption [29,30]. Nevertheless, there are no studies using MR on a broader range of dietary items due to the lack of single nucleotide polymorphisms (SNP) strongly associated with food consumption to be used as instrumental variables.

By virtue of the availability of the data from UK Biobank, we have recently been able to broaden the number of foods for which genetic instruments are available, identifying several causal food-health relationships [31]. We thus decided to use MR to investigate the causal effect of 40 foods/dietary patterns on 123 blood metabolites measured by nuclear magnetic resonance spectroscopy (NMR) available from a previous large GWAS by Kettunen *et al.* [32]. We detected 417 potentially causal links between food and metabolites, replicating previous findings and bringing novel insights, and discuss how these may be mediating the effect of food on health.

## Materials and Methods

To infer the causal relationships between food and metabolites we used Two-sample MR. In contrast to conventional MR, which would require instrument, exposure and outcome to be available as individual-level data for the same samples, the same analysis can be performed using summary statistics from genome-wide association studies. In this case, instead of directly estimating the effect of the SNP on the exposure and on the outcome, the parameter estimates from previous association studies for the two variables are used. MR can thus be performed, even if sample sizes of the two studies are different and there is no sample overlap - in the latter case the method is called Two-sample MR [33]. Apart from the obvious advantage of using the existing summary statistics, Two-sample MR minimizes the potential residual genetic confounding. We performed Two-sample MR using MR-base via R-package TwoSampleMR [34] (See detailed description: https://mrcieu.github.io/TwoSampleMR/).

For our study, exposure instruments were selected from the previous GWAS on food consumption conducted in UK Biobank data (up to N=445,799) [31] which included 25 traits for which valid instruments were available: consumption of beef, beer, bread, champagne/white wine, cheese, cooked vegetables, decaffeinated coffee, dried fruit, fresh fruit, ground coffee, instant coffee, lamb, non-oily fish, oily fish, pork, poultry, processed meat, red wine, salad, salt, spirits, tea, water adjusted for coffee, and vegetarianism and drink temperature.

In order to be able to distinguish between the independent effects of single food items and those arising due to the effect of dietary patterns, we defined 15 “Principal Component traits” (PC-traits) by first clustering the single food items using the iCLUST algorithm [35,36]. The defining and calculation of PC-traits is not a part of this paper and was done for our previous manuscript - therefore only a brief description is included here and more detailed information can be found in Pirastu *et al.* [31]. After clustering the single food items we split the resulting tree dendrogram into different layers depending on the items in each cluster and the degree of similarity. For example, Oily-Fish and Non-Oily fish were first grouped in an overall fish consumption variable *(Fish-PC1)* and then in a more general healthy foods measure together with *Fruit-PC1* and *Vegetables-PC1*. Finally, they were all used to estimate a measure of overall dietary pattern *(All-PC1-3).* Figure 1 shows the Sankey plot of the relationships between the different defined traits, and Figure 2 represents the loadings of each single food/drink on each of the main PC-traits (a full table of the PC-loadings can be viewed in Table S1 and is visualized in Figure S2). Higher values of *All-PC1* correspond to what is generally considered a “healthy diet”: higher consumption of vegetables, fruit and fish and lower consumption of meat, coffee and alcohol. *All-PC2* separates foods from drinks (alcohol containing beverages and coffee) with higher values corresponding to higher consumption of coffee and alcohol, and lower consumption of the rest of the foods. *Fruit-PC1* corresponds to higher consumption of dried and fresh fruit whereas higher value on *Psychoactive-PC1* corresponds to higher consumption of coffee and alcohol.

**Figure 1.**
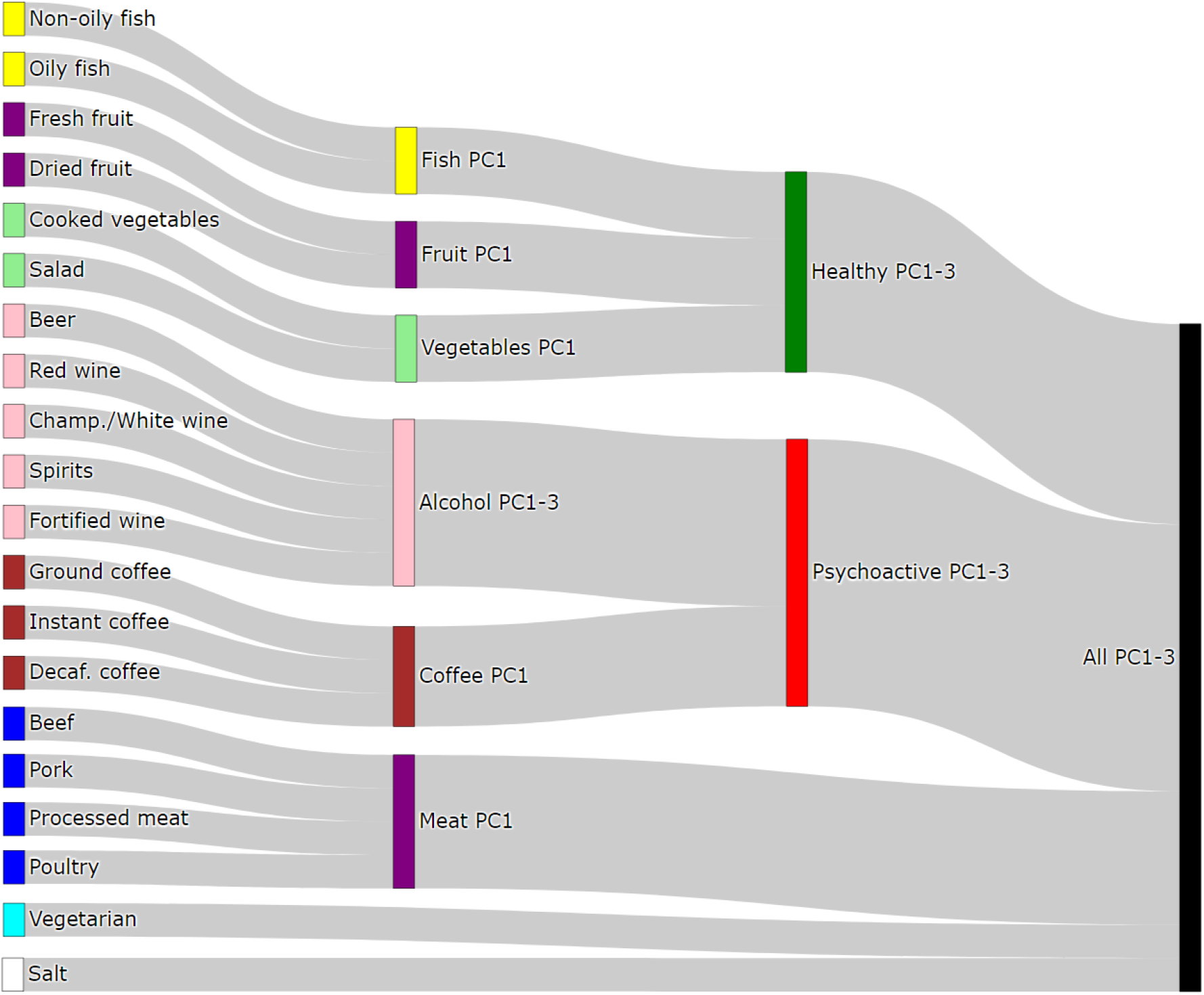
Sankey diagram of the relationships between dietary items and the principal components traits.

**Figure 2.**
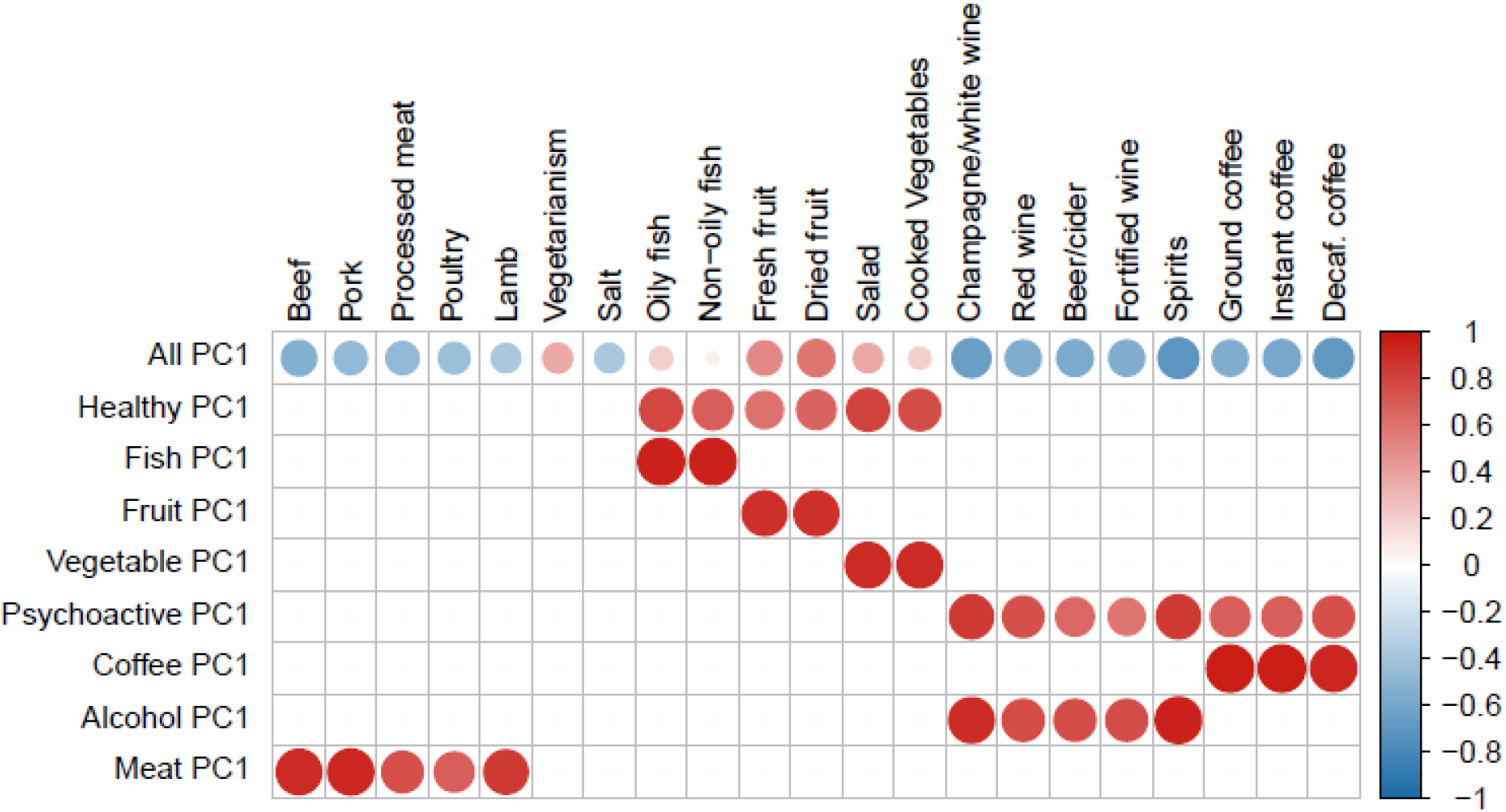
Loadings of PC traits. The plot represents the loadings of separate dietary items on each of the main PC-traits. Blank squares indicate that the corresponding item is not a component of the PC-trait. The size of the dots corresponds to the magnitude of the loadings whereas the color of the dots describe in addition the direction of the loadings.

The SNP effect estimates for the PC-traits were estimated by applying principal components analysis to the genetic correlation matrix of the items included in each group and by using the resulting rotation matrix to project the estimated effects of the SNPs on each of the items onto the PC space. Standard errors for the PC-traits effects were estimated using the single items SEs and the phenotypic correlation matrix.

As instruments we used independent (r2<0.001) SNPs significantly associated with the traits in the exposure GWAS (p-value < 5×10-8) for which the effect was not mediated through other confounders or health-related traits. In case of the PC-traits, SNPs were selected if they were associated (p-value <5e-8) with at least one of the items which participated in the trait definitions. Once extracted, we assigned to each selected SNP the lowest p-value amongst the traits of interest. Finally, we applied LD pruning (r<0.001) to the selected SNPs. We have previously shown that food frequency GWAS results are strongly affected by educational attainment (as a proxy of socioeconomic status) and by health-related traits such as body mass index, blood pressure and cholesterol, for which dietary advice is generally given [31]. This results in either bias by indication (where the behaviour is determined by health advice or belief, e.g. lower fat consumption in people with high cholesterol) or reporting bias (where people under- or over-report food consumption due to their health status, e.g. obese people under-report true fat consumption). If the biasing trait is heritable, this leads to spurious results in the GWAS, which can bias the MR results. In order to distinguish which variables are likely directly associated with the food of interest (and not mediated by health conditions) we have previously developed a method called Corrected to Uncorrected ratio filtering (CUR), which is based on the idea that if the SNP is directly associated with food preferences, then its effect should not change when adjusted for education status or health conditions [31]. In short, CUR corresponds to the ratio between the effect corrected for the aforementioned covariates and the uncorrected effect, and CUR substantially deviating from 1 is an indication of an invalid instrument. We thus selected only those SNPs which showed CUR = 1+/-0.05, which we have previously shown maximises the chances of selecting the correct SNPs. The SNPs effects for the outcomes were obtained from a previous GWAS on 123 plasma metabolites in 24,925 individuals [32].

As the main MR method we used the Inverse Variance Weighted method, with random effect standard error if heterogeneity p-value was less than 0.05/123. One of the problems of MR is when SNPs are associated with the outcome through causal paths which do not pass through the exposure of interest, also referred to as horizontal pleiotropy. In order to remove the SNPs with the highest heterogeneity (and thus likely are subject to horizontal pleiotropy) we used the method called MR-Radial [37].All analyses were thus run on the instruments selected with this method. As sensitivity analyses, we used MR-Median [38], MR-RAPS [39] and MR-Egger [40]. These methods have been thoroughly described elsewhere and are all sensitive to the breaking of different MR assumptions. When only one instrument was available the Wald ratio method was used. Finally, we defined as significant the food-metabolite relationships where Storey’s q-value [41] was less than 0.05. The analyses were run using R version 3.6.1 [42].

## Results

After correcting for multiple testing by using false discovery rate (FDR<0.05) via Storey’s q-values, 417 potentially causal relationships remained statistically significant. Most of these are associations related to atherogenic lipoproteins: very-low density lipoproteins (VLDL), intermediate density lipoproteins (IDL) and low-density lipoproteins (LDL), which all contain Apolipoprotein B (ApoB). Figures 3 and 4 are heatmaps reporting the relationships between food-items and -groups with the metabolites, Figure 3 depicts atherogenic lipoproteins and related metabolites, and Figure 4 depicts all other food-metabolite relationships of interest. For readability some metabolites which had only single significant associations are left out of these graphs. A full list of all significant food-metabolite relationships that we detected, is found in Table S2.

**Figure 3.**
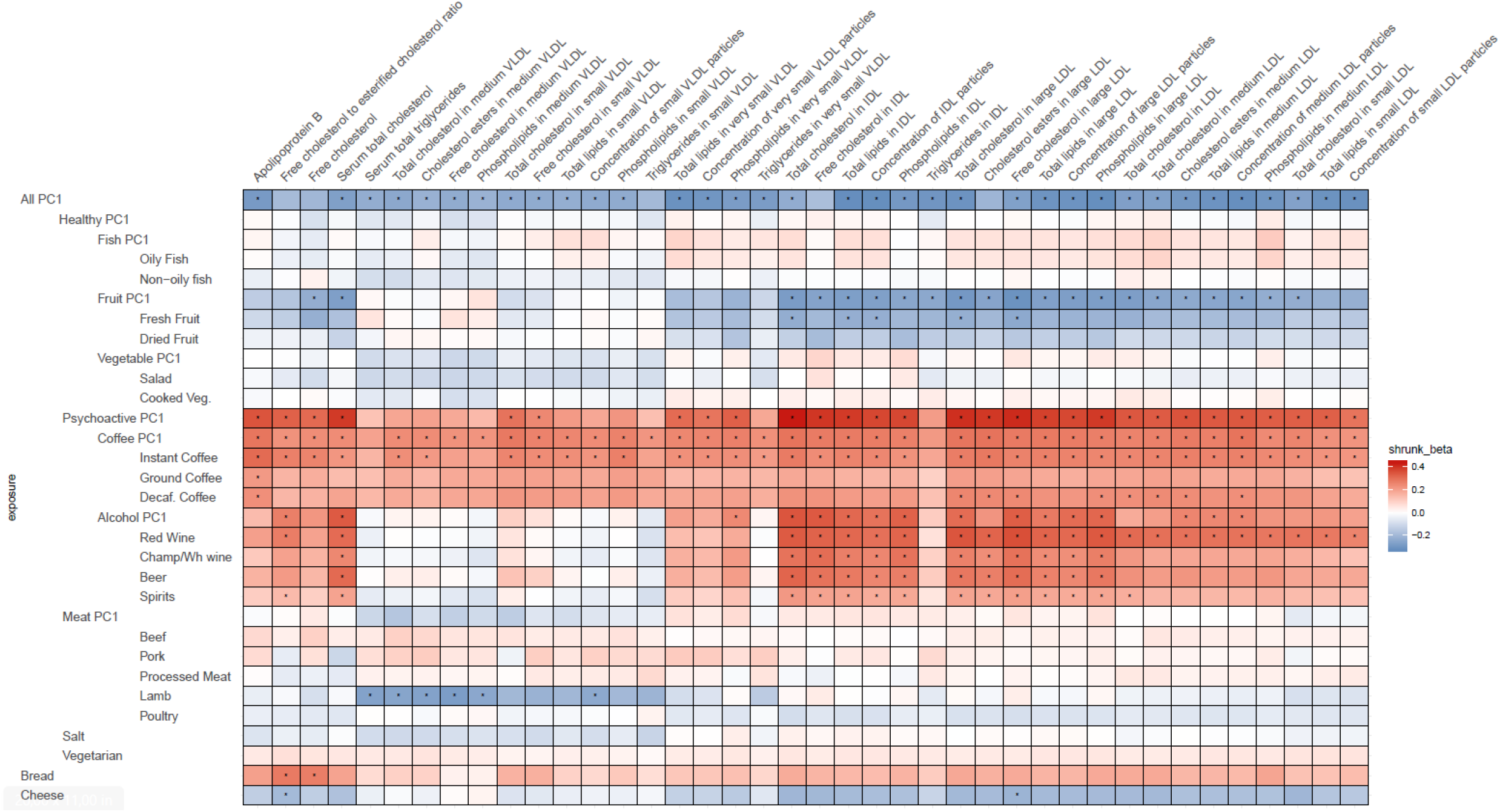
Heatmap of the relations of food traits with atherogenic lipoproteins and related metabolites (VLDL, IDL, LDL and related). Depicted are only the metabolites which showed significant association with the food-items or - groups. To facilitate meaningful visualisation and maximise the appearance of signal rather than noise, we applied a shrinkage method - imposing a bayesian prior assumption on the distribution of beta (mean 0, SD 0.1), and conjugating that with the likelihood of our results and then taking mean beta from the resulting distribution, thus shrinking estimates with larger SEs more towards 0. The color of the squares indicates the size and direction of betas after a shrinking procedure described in the Methods. The FDR-significant results are marked with “*” in the middle of the square.

**Figure 4.**
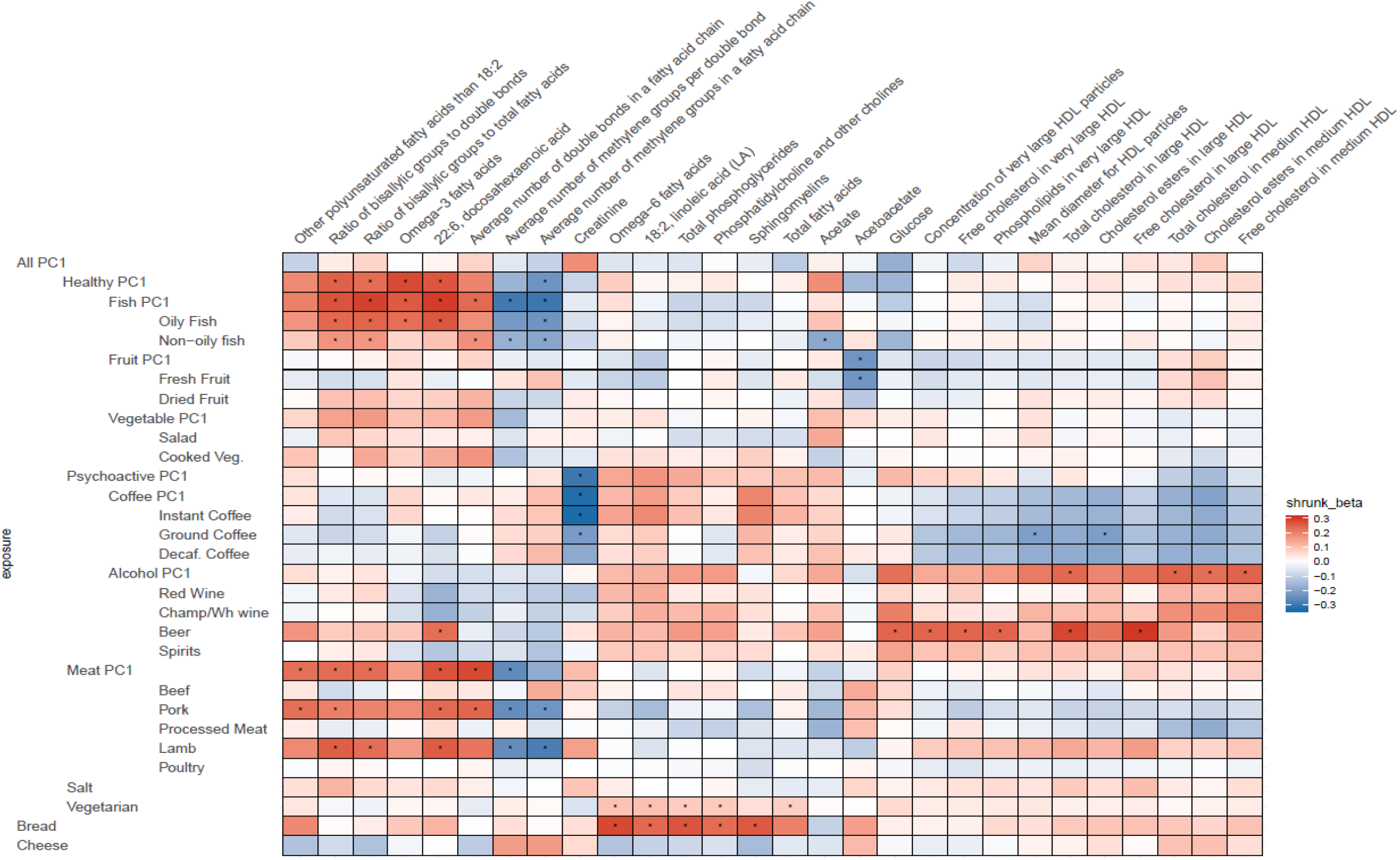
Heatmap of the relations of food traits and all other metabolites (not VLDL, IDL, LDL related). Depicted are only the metabolites which showed significant association with the food-items or -groups. To facilitate meaningful visualisation and maximise the appearance of signal rather than noise, we applied a shrinkage method - imposing a bayesian prior assumption on the distribution of beta (mean 0, SD 0.1), and conjugating that with the likelihood of our results and then taking mean beta from the resulting distribution, thus shrinking estimates with larger SEs more towards 0. The color of the squares indicates the size and direction of betas after a shrinking procedure described in the Methods. The FDR-significant results are marked with “*” in the middle of the square.

Out of the 417 significant results, 19 were based on the Wald ratio and 398 were based on the Inverse Variance Weighted method. For the latter, we used MR-Median, MR-RAPS and MR-Egger as sensitivity analyses to detect potential violation of assumptions. We found that the sensitivity analyses broadly agreed with the results of the main analysis with the exception of one case (*Alcohol-PC2* on “Description of average fatty acid chain length, not actual carbon number”), where the direction of the effect estimate from MR-Median was opposite to the one from the main analysis and thus this result cannot be considered reliable. In 16 cases (mostly when *Alcohol-PC2* was the exposure), the betas from MR-median were smaller (>50% effect difference) than the ones in the main analysis, which indicates that in these cases the effect sizes might be inflated. For some food-metabolite pairs, we detected some indication of heterogeneity. This was mostly related to PC-traits which reflect the effects of many different exposures and it is thus not unexpected.

Most significant relationships were found with the overall “healthy diet” trait (*All-PC1*), higher consumption of coffee (*Coffee-PC1*) and consumption of alcoholic beverages and coffee (*Psychoactive-PC1* and *-PC2*). The majority of the effects of these traits were on the same metabolites (VLDL, IDL and LDL related) and with opposite directions: namely showing negative correlation for *All-PC1*, where higher values correspond to a generally healthier diet, and positive correlations for *Coffee-PC1* and *Psychoactive-PC1.* These effects were very similar to each other in size, with betas ranging from -0.480 to -0.315 for *All-PC1,* from 0.327 to 0.436 for *Coffee-PC1* and from 0.426 to 0.778 for *Psychoactive-PC1*. The latter is likely partly due to the strong correlation between these lipid measurements.

*All-PC1* had significant associations with 43 of the measured 123 metabolites, which was, as expected, the highest number of associations we saw for any of the tested foods or PC’s. Interestingly these effects were mostly related to atherogenic lipoproteins: the components of VLDL, IDL and LDL of various sizes, and ApoB; whereas surprisingly there were no notable effects on any of the components of high density lipoprotein (HDL) particles or other metabolites measured (for example omega-3 or omega-6 fatty acids). Additionally, *All-PC1* had significant effects on serum total cholesterol and serum total triglycerides. When a significant causal relationship was found for a metabolite, the associations of this metabolite with each dietary item were examined. Often the effects on PC-traits were driven only by a subset of dietary items. For example, in the case of total lipids in IDL, although we see a clear effect of overall diet (*All-PC1*), it seems that this is driven primarily by the components of *Fruit-PC1* and *Psychoactive-PC1* while the remaining foods do not seem to play any role. This observation may explain some of the significant heterogeneity we detected.

Some strikingly clear patterns can be noted, when examining the results that are depicted on Figure 3 - for example looking at the significant positive effects that *Psychoactive-PC1* has on the components of IDL particles, large LDL particles, and serum total cholesterol. In the case of these results both subcategories - coffee and alcohol - clearly contribute to the overall effect of *Psychoactive-PC1*. More importantly, all the four subcategories of alcohol show independently significant effects. All the significant effects that *Psychoactive-PC1* has on the components of IDL particles and large LDL particles are accompanied by a contrary effect from *Fruit-PC1,* which shows significant negative effects. The latter indicates that higher consumption of fruits has a lowering effect on several IDL and large LDL components. In these cases, the effects of *Fruit-PC1* are with the same direction as the effects of *All-PC1,* indicating the healthier diet. Furthermore, this association pattern is also partly followed by the components of medium LDL particles, but in this case, when looking into the components of *Psychoactive-PC1,* alcohol seems to play less important role compared to coffee.

In most of the cases alcohol and coffee both contributed to the overall effect of *Psychoactive-PC1* and we cannot distinguish between the effect of the components (although alcohol seems to play a slightly larger role in the case of IDL particles). Therefore, in most of the cases *Alcohol-PC1, Coffee-PC1* and the items comprising these behave in the same way. Nevertheless, there is a group of clear and notable counterexamples: the components of medium VLDL and small VLDL particles. In these cases, *Coffee-PC1*shows clear positive effects and all the betas of all the coffee subgroups agree with the direction of these effects, whereas *Alcohol-PC1* and all of its subgroups show no clear effects and do not seem to be associated with the components of medium and small VLDL particles. Another interesting example that behaves differently from other items in the group of *Psychoactive-PC1* is beer, which has significant positive effects on glucose (β= 0.43, 95% CI: 0.15 ; 0.70), concentration of very large HDL particles (β= 0.47, 95% CI: 0.16 ; 0.77), free cholesterol in very large HDL (β= 0.45, 95% CI: 0.16 ; 0.75), phospholipids in very large HDL (β= 0.46, 95% CI: 0.16 ; 0.76), total cholesterol in large HDL (β= 0.53, 95% CI: 0.23 ; 0.83), free cholesterol in large HDL (β= 0.57, 95% CI: 0.27 ; 0.86), and on 22:6 docosahexaenoic acid (DHA, a subgroup of omega-3 fatty acids; β= 0.84, 95% CI: 0.32 ; 1.37). The latter is an example, where beer is clearly the odd-one-out compared to the effects of other alcohol subgroups, indicating that the effect comes from other ingredients in beer rather than alcohol.

The effects of *Meat-PC1* do not have much contribution from beef, processed meat and poultry and are driven mostly by pork (in the case of polyunsaturated fatty acids than other 18:2, and average number of double bonds in a fatty acid chain) or lamb (in the case of ratio of bisallylic groups to total fatty acids) or both (ratio of bisallylic groups to double bonds, DHA, and average number of methylene groups per double bond). Furthermore, it is noteworthy that lamb as a separate item had significant negative effects on several components of medium, large, very large and largest VLDL particles, namely on total cholesterol, cholesterol esters, free cholesterol and phospholipids in medium and large VLDL; total lipids and triglycerides in very large and largest VLDL; phospholipids in very large VLDL; concentration of small and large VLDL particles, and mean diameter for VLDL particles. Thus, lamb has significant negative effects on the components of those lipoproteins that are largest and with lowest density, whereas showing no notable effects on the components of any of the lipoproteins that are smaller and with higher density than small VLDL particles.

As expected, *Fish-PC1* had significant effects on omega-3 fatty acids and DHA and these effects were clearly driven by oily fish. Furthermore, looking at Figure 4, the results regarding creatinine notably stand out, namely there is a lowering effect of *Psychoactive-PC1* on creatinine, which is clearly driven by coffee (which in turn shows significant negative effect on creatinine). Surprisingly vegetarianism and bread share largely the structure of effects - they both have significant effects on omega-6 fatty acids (β= 3.73, 95% CI: 1.75 ; 5.70; β= 0.67 95% CI: 0.30 ; 1.04 respectively), 18:2 linoleic acid (LA, a subgroup of omega-6 fatty acids; β= 3.83, 95% CI: 1.86 ; 5.80; β= 0.55 95% CI: 0.18 ; 0.91), phosphatidylcholine and other cholines (β= 3.27, 95% CI: 1.31 ; 5.24; β= 0.53 95% CI: 0.17 ; 0.89) and total phosphoglycerides (β= 3.19, 95% CI: 1.23 ; 5.16;β= 0.63 95% CI: 0.25 ; 1.00). Of note, the results of vegetarianism are based only on one instrument and the method of Wald ratio was used.

## Discussion

In this study we have assessed the effect of long term exposure to single foods and food groups on blood metabolite profiles using Mendelian randomization. We have in general found that in many cases these changes are not due to specific food items but are related to general dietary patterns. This could be due to the fact that most of the metabolites assayed are linked to lipid profile and it does not exclude more specific biomarkers being discovered for single items.

Some of the significant results that we found between healthier diet (represented by higher values on *All-PC1)* and metabolites (such as total lipids in large VLDL, small VLDL, large LDL, medium LDL, small LDL and IDL) replicated the findings of a recent observational study investigating association between metabolites and healthy diet [43], while for others (ApoB, LDL cholesterol (LDL-C), IDL cholesterol (IDL-C), serum total cholesterol, total lipids in very small VLDL) they showed a non-significant association with the same direction. Many of the results that we found elaborate the lipid profiles of lipoprotein subclasses in more detail than the previous studies. Our results regarding the effects of alcohol on IDL and large LDL related lipids conflicted with some observational studies’ findings [18,44], but agreed with another MR-study [45], whereas in our study we showed that the same relationships hold for each of the alcohol subgroups as well. Our finding of vegetarianism raising 18:2 linoleic acid replicated a previous finding from a randomized trial conducted on subjects with T2D [46]. Furthermore the finding of vegetarianism raising omega-6 replicated a previous observational finding [47]. We also saw a significant positive effect of oily fish on omega-3 fatty acids and DHA, which are well known causal relationships [48]. The fact that our results conflicted with some observational findings, but aligned with known causal relationships and with a randomized trial, highlights the strength of MR studies as an intermediate step between observational studies and clinical trials and validates the utility of our approach.

One of the most notable patterns in our results was the one regarding IDL particles and large LDL particles, where higher consumption of coffee and alcohol had elevating effects, whereas higher consumption of fruits and higher value on *All-PC1* (i.e. healthier overall diet) had lowering effects. Higher levels of atherogenic particles and their components is part of a less desirable blood lipoprotein profile due to elevated risk for several cardiovascular diseases [49]. For example, IDL-C has been shown to be associated with the degree and frequency of CAD independent of LDL-C [50]. Furthermore, different properties of LDL, like size, the amount of cholesterol esters and cholesterol, and fatty acid composition are all considered to be aspects of its CVD-causing capability [51]. Large LDL particles have been found to be predictive of coronary artery disease in numerous studies, also when accompanied by low or normal triglyceride levels [52]. IDL and LDL may have slightly different mechanisms by which they advance the progress of coronary artery disease, while still having a direct association with CVD [52]. The clear pattern of associations we saw on IDL and large LDL indicate that these metabolites are likely affected by dietary habits. Since these metabolites are largely shown to affect cardiovascular health, and Mendelian randomization results indicate possible causal pathways, there is considerable scope for further investigation of these results. Furthermore, our results indicate that the harmful effects alcohol and coffee have on cardiovascular health are at least partly mediated by IDL and large LDL lipoproteins.

On the other hand, as we saw from the results, in addition to harmful effects, alcoholic beverages can have somewhat beneficial effects as well, namely elevating effects on the components of large and medium HDL. The effects on large HDL were largely driven by beer or seen only for the beer subgroup. Nevertheless, it is important to note that even though there is epidemiological evidence of HDL having a beneficial role for cardiovascular health, the extent and specific mechanism is still not fully understood or proven [53], and the studies have failed to show a causal link [54]. Furthermore, there was a surprising positive effect of beer on DHA, which has been shown to have a cardioprotective effect, as concluded in a recent large meta-analysis of randomized control trials [55]. This effect on DHA was in the opposite direction compared to other alcoholic beverages, indicating that the beneficial effect is likely not due to alcohol itself but some other ingredient in beer. We propose that the beer-DHA relationship is worth further investigation and it would be especially interesting to compare regular beer with non-alcoholic beer to detect whether the beneficial effects remain. Furthermore, even if alcoholic beverages can have some beneficial effects, one can not underestimate the negative impact of elevated IDL and LDL levels on health coming from alcohol itself.

Many of our significant results were related to an overall healthier diet (*All-PC1*), which seems to affect mostly LDL, VLDL, IDL and related subclasses. The fact that the effect sizes were very similar, can be partly explained by the strong correlations between the different metabolites explained in the results section. Another explanation might be the effect that *All-PC1* has on ApoB. Namely, healthier values on *All-PC1* have a lowering effect on ApoB. The latter is the protein part of all the lipoproteins formed in the liver that carry cholesterol and triglycerides to the cells of the body. Those lipoproteins are VLDL, IDL, LDL. As there is only one ApoB protein per each such lipoprotein [56], one can assume that the total amount of ApoB will reflect the whole amount of atherogenic lipoprotein particles. This is important, because it shows that ApoB, being a component of all atherogenic lipoprotein particles, might reflect the actual CVD risk better than the amount of cholesterol in any lipoprotein particle type on their own [57]. The latter has been shown in some studies, for example Walldius and Jungner [58] found that ApoB levels predicted the risk of CVD better than LDL-C and especially in patients who didn’t have elevated LDL-C levels. Ference *et al.* [59] came to the conclusion that the risk of CVD posed by LDL particles is more determined by the concentration of LDL particles measured by ApoB compared to the mass of cholesterol in LDL particles. Because of the lowering effect of *All-PC1* on ApoB, we can assume that ApoB might be one of factors playing a substantial role in the relationship between diet and health, and that healthy diet changes that contribute to lowering ApoB levels might have the potential to actually reduce the risk of CVD.

We saw a significant increasing effect from *Coffee-PC1* on several VLDL, IDL and LDL lipoprotein subclasses and their components and in addition a negative effect on the mean HDL diameter and cholesterol esters in large HDL. Effects on other HDL parameters were not significant, but there is a notable trend towards a negative correlation. Overall, a higher value on *Coffee-PC1* results in a more unhealthy lipid profile, raising ApoB, serum total cholesterol, VLDL, IDL and LDL levels and their constituents. Nevertheless, in the case of coffee, the results regarding cardiovascular health are controversial and mostly show either beneficial or neutral effects [60] or that moderate consumption is unlikely to have adverse effects [61]. A recent review article concluded that filtered coffee consumption up to 6 cups a day did not increase the risk of cardiovascular diseases, but in fact moderate coffee consumption has been associated with a reduced disease risk [62]. Because coffee contains various biologically active substances besides caffeine, it is hard to pinpoint the different effects we see, compared to the ones in previous studies, to any specific compound. Raised ApoB levels indicate that the overall atherogenic particle amount is higher, which, as mentioned before, is a good predictor of CVD risk. Raised VLDL levels might be due to cafestol, a common ingredient in coffee, having an effect of increased VLDL particle assembly rate in the liver [63]. Raised serum cholesterol and LDL levels associated with coffee consumption have been documented before and are in line with our results [64,65]. Overall, we did not see any beneficial effect of coffee on lipid profile in our results, in fact coffee turned out to have a negative effect on a larger variety of atherogenic lipoproteins than other dietary items. Hence, the total effect of coffee on health remains still unclear, but we propose that the harmful effect of coffee on health might be mediated by ApoB and thus via VLDL, IDL and LDL.

We saw significant increasing effects of vegetarianism on omega-6 fatty acids and on its subgroup 18:2 linoleic acid. The omega-6 result replicated a previous finding by Kornsteiner, Singer and Elmadfa [47], however our MR-analysis showed the potential for a causal relationship. The effect of vegetarian diet on 18:2 linoleic acid replicated a previous result of a randomized controlled trial in T2D patients [46], whereas our results show that this finding applies for the general population in a larger sample as well. These results are important in the context of omega-6/omega-3 ratio, since too high omega-6/omega-3 ratio is associated with multiple diseases, including cardiovascular disease, cancer, inflammatory and autoimmune diseases [66]. The recommended ratio between omega-6 and omega-3 is about 4:1 [66], but vegetarian diet has been shown to have a ratio of 10:1 [47]. Thus, a higher amount of omega-6 may be driving the ratio to an undesired direction and consequently the individuals on a vegetarian diet might, in addition to adding omega-3 rich foods to their menu, need to also pay extra care in reducing the intake of omega-6 rich foods in order to maintain a healthy omega-3/omega-6 ratio.

Our study has several limitations: for some items only a few SNPs were available to use as an instrument in MR; self-reported dietary data is a difficult trait to investigate since it encompasses several biases - we aimed to mitigate this issue by using Corrected-to-Uncorrected Ratio [31]; MR analysis can suffer from horizontal pleiotropy, we tried to mitigate this issue by using sensitivity analyses. Nevertheless, our study has several strengths: to our best knowledge, we are the first ones to perform MR analysis between dietary items and blood metabolites with such a large amount of dietary SNP instruments available; we found several distinct patterns that shed more light on how dietary changes might affect cardiovascular health; we found multiple interesting associations that are worth further investigation via feeding studies or randomized trials. We investigated the effects on metabolites profiled with NMR spectroscopy, which encompass mostly lipid profiles. Future studies with proteomics and metabolite data from mass spectrometry might give more detailed insight about the mechanisms by which food affects health.

In conclusion, we aimed to investigate the relationships between dietary items and blood metabolites in order to gain more insight into the mechanisms by which food affects health. Mendelian randomization proved a useful method for fulfilling this aim. Moreover, we replicated several previous findings and known associations, which validates the method used. We did not detect any reported causal relationships in the literature conflicting with our results, however occasionally our results conflicted with previous observational studies. This demonstrates the strength of MR studies and indicates that some of the previously reported findings might be confounded through other unobserved factors. Nevertheless, in order to give actual dietary intervention suggestions, additional thorough investigations should be carried out via feeding studies or randomized trials. We believe that many of the potentially causal relationships that have been described here, have promising potential for further investigation.

## Supplementary Materials

**Figure S1**: Loadings of all principal component traits (The plot represents the loadings of separate dietary items on each of the PC-traits. Blank squares indicate that the corresponding item is not a component of the PC-trait. The size of the dots corresponds to the magnitude of the loadings whereas the color of the dots describe in addition the direction of the loadings.), **Table S1**: Loadings of all separate dietary items on each of the principal component traits., **Table S2**: All FDR-significant results between dietary traits and blood metabolites.

