## Supplementary figures and images for "Mendelian randomization identifies the potential causal impact of dietary patterns on circulating blood metabolites"

### Figure S1

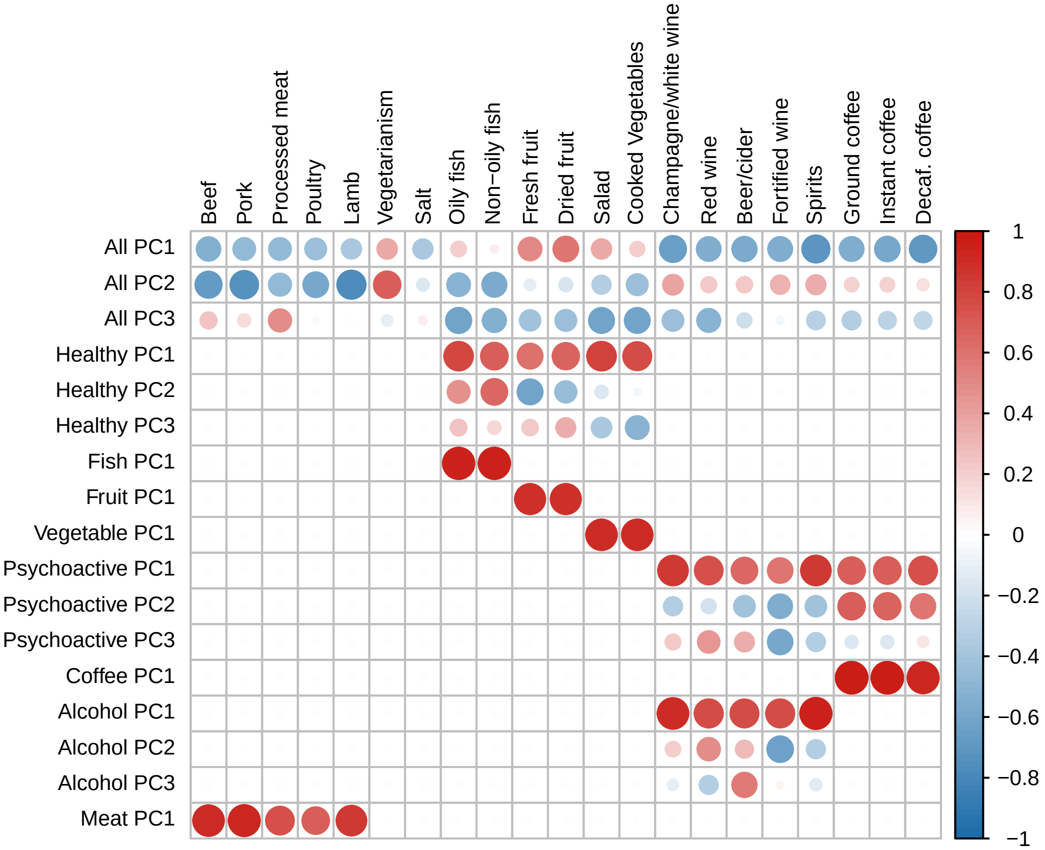
